# Implicit representation of viruses fails to capture impacts of virus-induced mortality in global marine ecosystem models

**DOI:** 10.64898/2026.07.21.739416

**Authors:** David Talmy, Eric Carr, Paul Frémont, David Demory, Christopher L. Follett, Oliver Jahn, Daniel Muratore, Stephen J. Beckett, Debbie Lindell, Joshua S. Weitz, Stephanie Dutkiewicz

## Abstract

Virus-induced mortality influences plankton biogeography, community structure, and ocean elemental cycles. However, quantification of virus-induced impacts remains challenging and often limited in scope. An alternative to explicit inclusion of viral dynamics in biogeochemical models is to represent viral effects implicitly, by assuming that mortality increases quadratically with cell or biomass density. Using 1D and 3D configurations of a nutrient-phytoplankton-zooplankton-virus-detritus (NPZVD) model, we ask whether the implicit quadratic mortality assumption captures patterns of virus-induced mortality, and its impact on biomass and primary production. The 1D water-column configuration shows that, at the onset of the spring bloom, the quadratic, implicit representation imposes viral losses on phytoplankton density instantaneously, which limits spring bloom formation. This is in contrast to the explicit representation, which allows initial bloom formation to proceed unhampered initially, but imposes a far stronger viral mortality later in the year driven by high rates of host-virus contact due to high phytoplankton and viral densities that take time to accumulate. By comparison to the implicit model, explicit resolution of viruses within the 3D global model shows strong potential for viruses to prematurely terminate phytoplankton blooms. Biogeochemical models would therefore benefit from explicit representation of viral infection insofar as models can be developed that adequately recapitulate *in situ* observations.

**Key Points:**

- Global model reveals significant spatial heterogeneity arising from explicit, rather than implicit, representation of viruses
- Implicit representation of viral dynamics fails to capture the potential for viruses to terminate phytoplankton blooms
- Biogeochemical models require explicit representation of viruses to adequately simulate their effect on marine systems

## 1 Introduction

By cycling nutrients between inorganic and organic forms, ocean microbes influence global ecosystems and biogeochemical cycles. Viruses are numerically abundant microbial community members throughout the global ocean, typically exceeding bacterial abundances in the surface ocean by 1x to 100x (Wigington et al., 2016). By infecting and lysing microbial hosts, viruses have been invoked as terminators of phytoplankton blooms (Lehahn et al., 2014), and are responsible for an appreciable but highly variably and poorly constrained proportion of microbial mortality. When viruses lyse their hosts, they convert organic molecules and nutrients within cells to new virus particles and detritus. Articulated in the viral shunt hypothesis (Fuhrman, 1999; Wilhelm & Suttle, 1999), material resulting from viral lysis may be ‘shunted’ into dissolved organic matter, and retained in the surface where it may be consumed or remineralized by bacteria or phytoplankton. Alternatively, the viral shuttle hypothesizes that viruses promote carbon removal from the surface ocean since lysed material may contain sticky polysaccharides that cause particle aggregation and the formation of marine snow (Weinbauer, 2004; Sullivan et al., 2017).

Zooplankton grazing is the process thought to be responsible for the majority of phytoplankton mortality (Calbet & Landry, 2004) and as such, zooplankton are usually resolved explicitly within biogeochemical models. The potential for viruses to influence elemental cycling warrants their inclusion in models, yet in practice, including viruses in large-scale ocean models presents a series of challenges. Viruses are extremely diverse, and each virus can infect only a subset of plankton hosts. Viruses spend a considerable portion of their life-cycle inside host cells, and the molecular processes underlying diversity in when and how viruses lyse their hosts are not fully understood (Edwards, 2018; Silveira et al., 2021). Nevertheless, the impact of viruses on ocean ecology and nutrient cycling is beginning to be evaluated in models (Weitz et al., 2015; Weitz, 2016; Frémont et al., 2026a). For example, an empirical model evaluated the impact of viruses on phytoplankton mortality and carbon dynamics in the California Current Ecosystem (Talmy et al., 2019), dynamical modeling has been used to explain biogeography of *Micromonas* and associated viruses (Demory et al., 2021), and a three-dimensional model has simulated the dynamics of *Emiliania huxleyi* viruses in the North Atlantic (Nissimov et al., 2019). Nevertheless, inclusion of the full range of ecologically dominant viruses within models presents a significant challenge, preventing robust quantitative assessment of their biogeochemical and ecological influences on marine ecosystems.

An alternative to explicit inclusion of viruses in models, is to simulate their ecological and biogeochemical impacts implicitly (Figure 1). Implicit inclusion of viruses is typically done by assuming that viruses are responsible for density-dependent host mortality following a quadratic function of host biomass density (Stock et al., 2014; Behrenfeld & Boss, 2014). The quadratic function is invoked in situations where other loss processes, such as zooplankton grazing or exudation, cannot reasonably be assumed to fully constrain phytoplankton or bacteria biomass density. However, to the best of our knowledge, the assumption that virus-induced mortality increases with the square of biomass density has not been directly tested with observational data, and the effects of the quadratic mortality function has not been shown to mimic the impact of viral infection on ecosystem structure and function.

**Figure 1:**
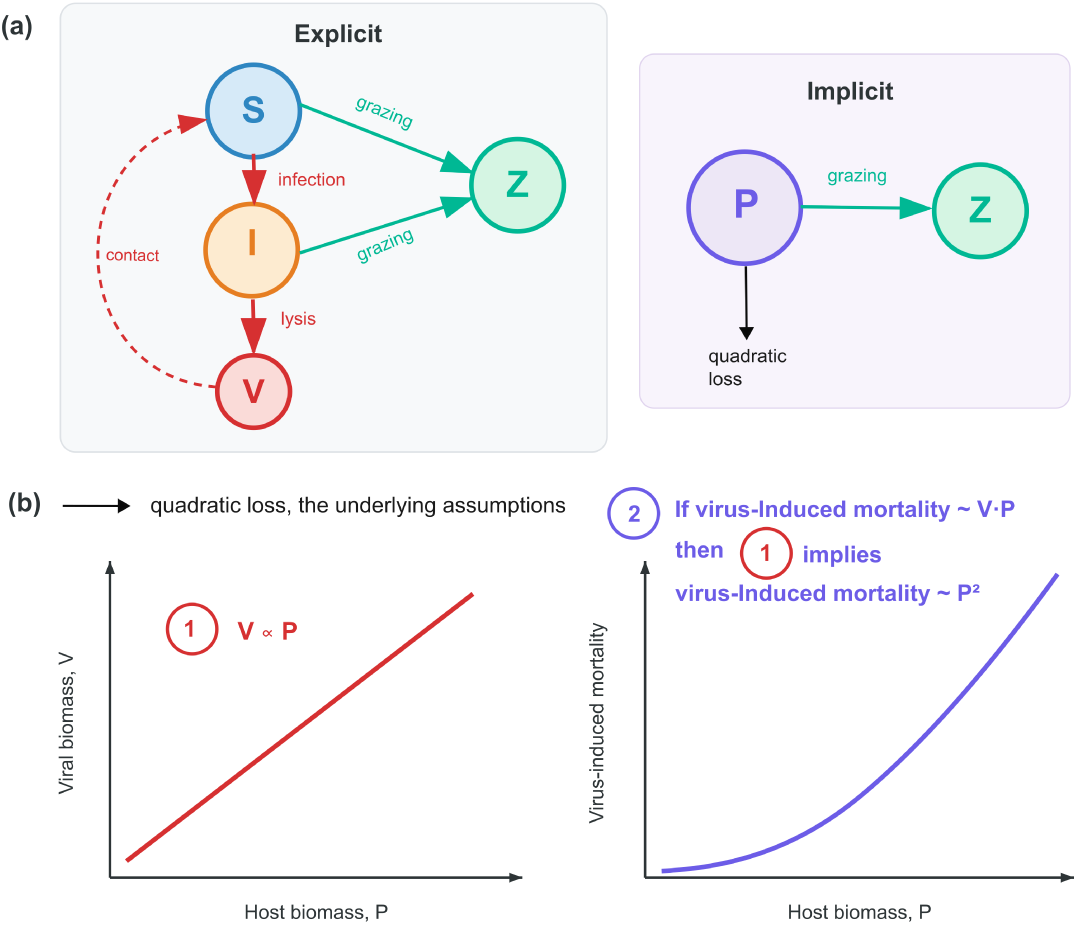
Schematic representation of explicit and implicit representations of viral infection within vDarwin (a), and a representation of the assumptions underlying the implicit representation (b). Explicit viral infection is described in Equations 2 to 5 and Frémont et al. (2026b). The implicit representation assumes that mortality increases with the square of biomass density.

A range of methods exist for *in situ* quantification of virus-induced mortality. Arguably the most widely applied is the modified dilution technique (Kimmance et al., 2007) which has the benefit of quantifying mortality of the entire microbial community (Pasulka et al., 2015), but suffers numerous limitations associated with artifacts arising from incubation of diverse communities in bottles. Alternative approaches infer viral mortality by identification of viruses in electron microscopic images of host cells prior to lysis (Proctor & Fuhrman, 1990). More recently, single-cell techniques have been developed that quantify molecular signatures of infection in individual cells (Ku et al., 2020; Mruwat et al., 2021; Brüwer et al., 2024). When combined with virus life-history traits (e.g. adsorption rates, burst size, etc.) in ecologically representative organisms such as *Prochlorococcus*, these molecular data can be used to infer rates of virus-induced mortality (Mruwat et al., 2021; Carlson et al., 2022; Beckett et al., 2024). This explicit link between life-history traits and molecular markers of infection is a natural fit with trait-based models such as the MITgcm-Darwin system (Follows et al., 2007; Dutkiewicz et al., 2020), and have recently been leveraged to initiate a new version of a virus-explicit model - vDarwin (Frémont et al., 2026b).

Frémont et al. (2026b) explored the sensitivity of ocean productivity and trophic conditions to viral dynamics. vDarwin represents a novel opportunity to ask whether explicit inclusion of viruses is necessary, or if their effects can be captured implicitly with the quadratic mortality approach. Here, we first ask whether explicit representation of virus-induced mortality in an ecosystem model follows a pattern consistent with the quadratic closure assumption. We then ask whether quadratic closure can mimic the effects of viruses on primary production and plankton biomass. Overall, we find that the quadratic closure does not recapitulate the dynamics of mortality, biomass and productivity that emerge when viruses are resolved explicitly. We discuss the implications of our findings for the broader goal of including viruses in ecosystem models.

## 2 Methods

We first provide an overview of the vDarwin-MITgcm and the descriptions of explicit and implicit viral infection. We then describe tests that were used to compare the impact of explicit and implicit viral infection on ecosystem dynamics.

### 2.1 Overview of vDarwin-MITgcm

The Darwin-MITgcm simulates a set of biogeochemical tracers, *X* (mmol C m*^−^*^3^), representing phytoplankton, zooplankton, inorganic nutrients and detrital material (Follows et al., 2007). Each biogeochemical tracer *X* is transported and subjected to biological sources and sinks following a mass-balance equation:

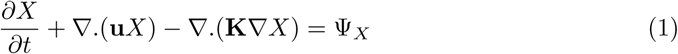

where ∇ is the spatial gradient operator, **u** is the three dimensional velocity vector, **K** are the mixing coefficients and Ψ*_X_* are the biological sources and sinks of *X* (e.g. nutrient uptake, grazing, etc.).

#### 2.1.1 Modeling virus-induced mortality

Transfer of material between tracers due to nutrient uptake and grazing follow the same equations used in the Darwin-MITgcm (Dutkiewicz et al., 2015). Following Frémont et al. (2026b), viral dynamics were added as follows. Let *S* represent the biomass density (mmol m*^−^*^3^) of a phytoplankton that is susceptible to viral infection, and let *I* (also mmol m*^−^*^3^) represent phytoplankton biomass that is infected by a virus. The total phytoplankton biomass *P* is then *S*+*I*. Sources and sinks Ψ*_S_* (mmol m*^−^*^3^ day*^−^*^1^) of the susceptible type are associated with nutrient uptake, abiotically induced mortality (e.g. mortality associated with reactive oxygen species (Ma et al., 2018)), viral lysis, and grazing, such that:

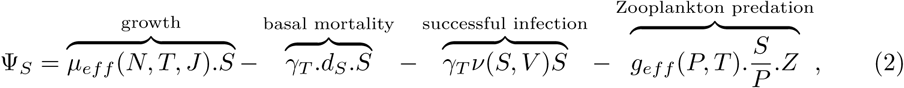

where *µ_eff_* is the effective growth rate (day*^−^*^1^), *d_S_* is the mortality rate (day*^−^*^1^), *g_eff_* is the effective grazing rate (day*^−^*^1^), *T* is the temperature (*K*), *N* is the limiting nutrient concentration (mmol m*^−^*^3^), *J* is irradiance (*µ*mol photons m*^−^*^2^*s^−^*^1^) and *γ_T_* is a temperature modulation coefficient (Dutkiewicz et al., 2020). The rate of virally mediated removal of susceptible biomass *ν* is modified for the explicit and implicit representations of viral infection, as specified in the following sections.

#### 2.1.2 Explicit viral infection

Here, if *V* is extracellular virus biomass density (mmol m*^−^*^3^), removal of biomass from the susceptible class follows the law of mass action, representing contact-dependent infection:

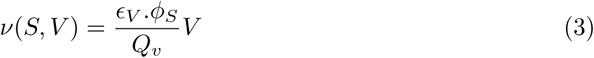

where *ɛ_V_* is the probability of a successful infection (non-dimensional), *ϕ_S_* is the adsorption rate of the virus (m^3^ day*^−^*^1^), and *Q_v_* represents the virus carbon quota (mmol C virus*^−^*^1^). Infected cells are then transferred to an infected biomass class *I*, with the following sources and sinks:

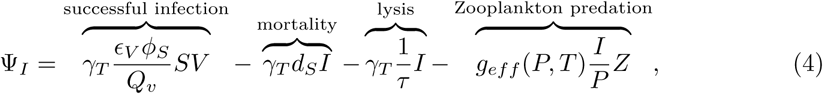

where *τ* is a timescale of infection (day), equivalent to the inverse of the lysis rate. Source and sinks of the extracellular virus are then:

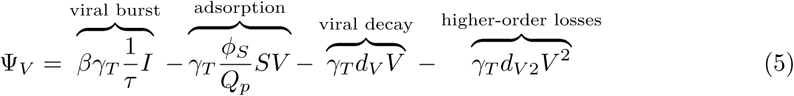

where *β* is the burst size (virions per cell), *Q_p_* is the phytoplankton carbon quota (mmol C cell*^−^*^1^), *d_V_* is the viral decay rate (day*^−^*^1^), and *d_V_*_2_ is the quadratic mortality of the virus (m^3^ day*^−^*^1^ (mmol C)*^−^*^1^). The viral decay accounts for abiotic degradation (Wilhelm et al., 1998) and quadratic mortality represents particle attachment (Yamada et al., 2020) and grazing by bacteria and protists (Suttle & Chen, 1992).

#### 2.1.3 Implicit viral infection

Arguably the simplest representation of virus-induced mortality assumes that the number of cells (or amount of biomass) lost due to viral infection is linearly proportional to the abundance (or biomass density) of both hosts and their viruses. In other words, virus-induced mortality ∼ *P* · *V*, where *P* and *V* are phytoplankton and virus abundance or biomass density, respectively. However, even this simple representation requires knowledge of the viral abundance, which is unknown in models that do not explicitly represent viral dynamics. A simplifying assumption is to assume that viral abundance is linearly proportional to host abundance, i.e. *P* ∼ *V*. In this case, the assumption that virus-induced mortality scales linearly with host and virus abundance is equivalent to assuming that mortality scales with the square of host abundance (Figure 1b).

Our description of implicit virus-induced mortality within the vDarwin-MITgcm is a result of the assumptions articulated in Figure 1b). Here, the *rate* of virus-induced mortality (as opposed to the flux, *ν*) increases linearly with the biomass of the susceptible type, i.e. *ν*(*S, V*) = *δ_vv_S*. We made different assumptions regarding the value of *δ_vv_*, which are described below.

### 2.2 Ecological and physical configurations

For simplicity, the configuration of vDarwin that we use here has a single phytoplankton type infected by a single virus with life-history traits and infection dynamics consistent with *Prochlorococcus* and their cyanophage (though note that vDarwin has been written to include multiple hosts and viruses). Model parameters associated with viral infection are described in Table 1. The full model was solved in one-dimensional and three-dimensional configurations. In the 3D case, the hydrostatic primitive equations are solved under the Boussinesq approximation (Marshall et al., 1997), and circulation is constrained by altimetric and hydrographic observations through the ECCO-GODAE state estimates (Wunsch & Heimbach, 2007). In the 1D configuration lateral advection and diffusion are neglected, and Equation 1 retains only vertical terms that are configured with hydrographic profiles observed at the Bermuda Atlantic Time-Series site (Wu et al., 2021).

**Table 1:**
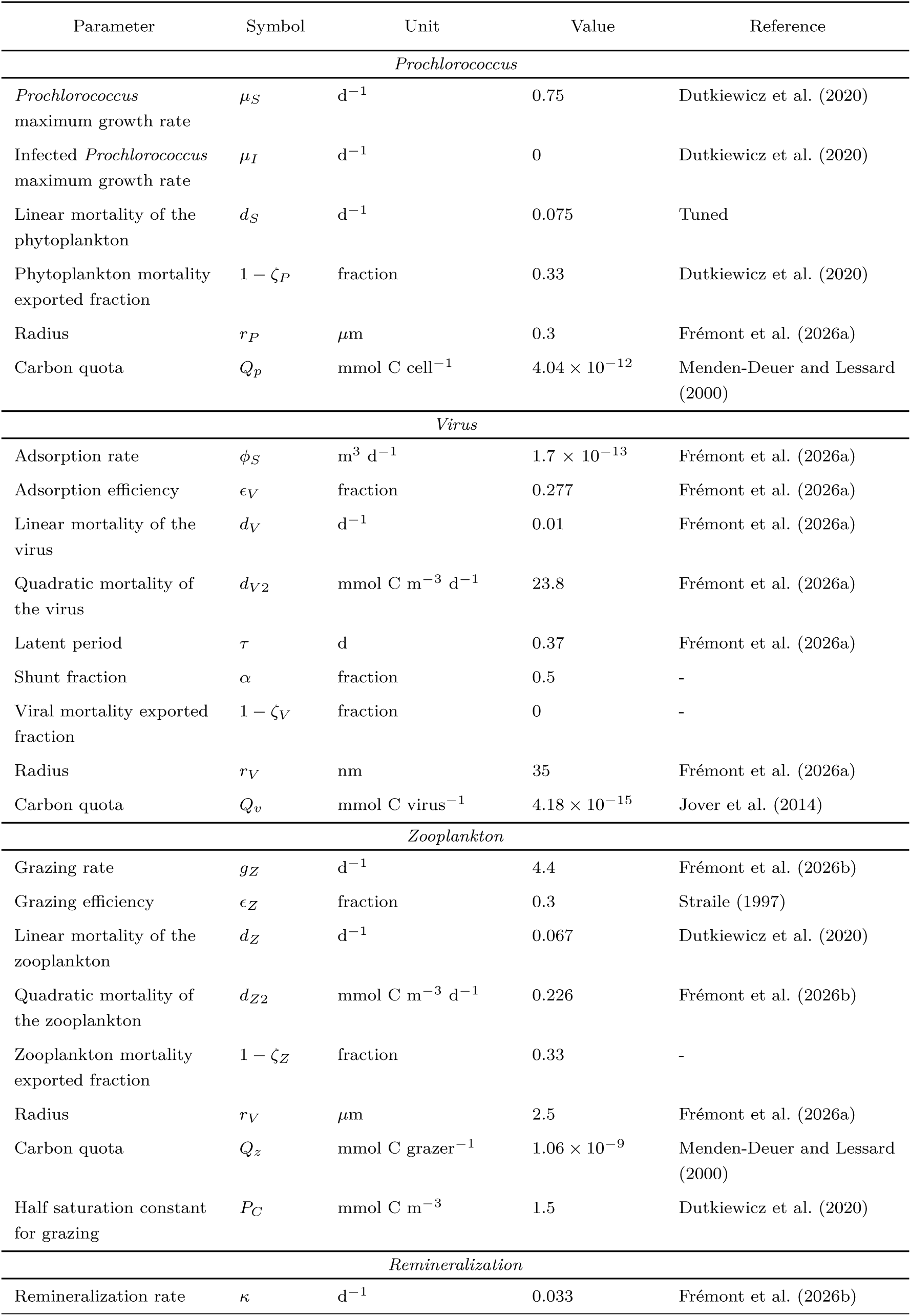
Model parameters used in the current configuration of vDarwin. Additional parameters specifiying rates of nutrient uptake and photosynthetic affinity are specified in Dutkiewicz et al. (2015) and Frémont et al. (2026b).

### 2.3 Tests and sensitivity analyses

We ran 1D and 3D simulations with the explicit virus, using parameter values from Frémont et al. (2026a, 2026b) that were tuned for the *Prochlorococcus*-cyanophage system. Parameter values are specified in Table 1. Fifty per cent of lysed material was directed to particulate and dissolved organic material, respectively, representing a 50-50 shunt-shuttle partition, consistent with viruses exerting minimal influence on net primary production relative to virus-free control simulations (Frémont et al., 2026b). To simplify the model and isolate the role of viral dynamics, the temperature coefficient *γ_T_* was set to 1 in all simulations, including all equations that constrain phytoplankton and zooplankton growth and mortality. We used these simulations with explicit viral dynamics as a basis to ask two related questions: First, how well does the quadratic function explain the relationship between virus-induced mortality and host biomass? Second, is the implicit model able to capture the imprint of viral infection on ecosystem processes?

To address the first question, the explicit model was treated as ‘truth’, and the value of *δ_vv_* was chosen such that the quadratic function gave the best possible fit (with the SciPy v1.16.3 curve fit function using default optimizer settings) to the relationship between virus-induced mortality (defined as *γ_T_*^1^ *I* in Equation 4) and host biomass, *S*, that emerged in the explicit model. In this first case, no simulations with vDarwin-MITgcm were conducted with the implicit representation. Instead we used the explicit model output to ask in a post-hoc manner whether the quadratic closure representation mimics the relationship between virus-induced mortality and host biomass.

We then conducted sets of simulations in 1D and 3D configurations of vDarwin-MITgcm assuming different, fixed values of *δ_vv_*, with implicit viral lysis again directed to a 50-50 shunt-shuttle partition and no temperature sensitivity of biological processes. The goal here was to search for values of *δ_vv_* that allow the implicit model to capture spatio-temporal variation in biogeochemical properties such as primary productivity, phytoplankton carbon density and export, without explicitly modeling host-virus dynamics. In doing so, we tested the ability of the implicit model to capture viral imprints on key ecosystem processes and states.

## 3 Results

We first describe the ability of the implicit model to capture the relationship between virus-induced mortality and host biomass in a one-dimensional configuration of the vDarwin-MITgcm. Using the same 1D configuration, we evaluate the effect of the explicit and implicit model on predictions of plankton biomass and productivity. In our comparisons, we first assume a single, fitted value of the quadratic coefficient in the implicit case, but we go on to conduct sensitivity studies to explore the ability of the implicit model to capture ecosystem dynamics across a range of assumed values of the quadratic coefficient, *δ_vv_*. Equivalent results are then reported for three-dimensional simulations.

### 3.1 Implicit vs. explicit descriptions of virus-induced mortality

Recall (Figure 1b) that an underlying assumption of the implicit model is a linear (i.e. proportional) relationship between virus and host biomass density. In Figure 2a we show that a linear relationship (green line) cannot capture the relationship that emerges between these two variables when viruses are modeled explicitly (blue dots in Figure 2; note the negative R^2^). The dynamic relationship between virus and host biomass density leads to qualitatively similar patterns in the relationship between virus-induced mortality and host biomass density (Figure 1b). As a result, the quadratic representation (green line, Figure 2b) can explain at most 6% of the relationship between virus-induced mortality and host biomass density when viruses are explicitly modeled.

**Figure 2:**
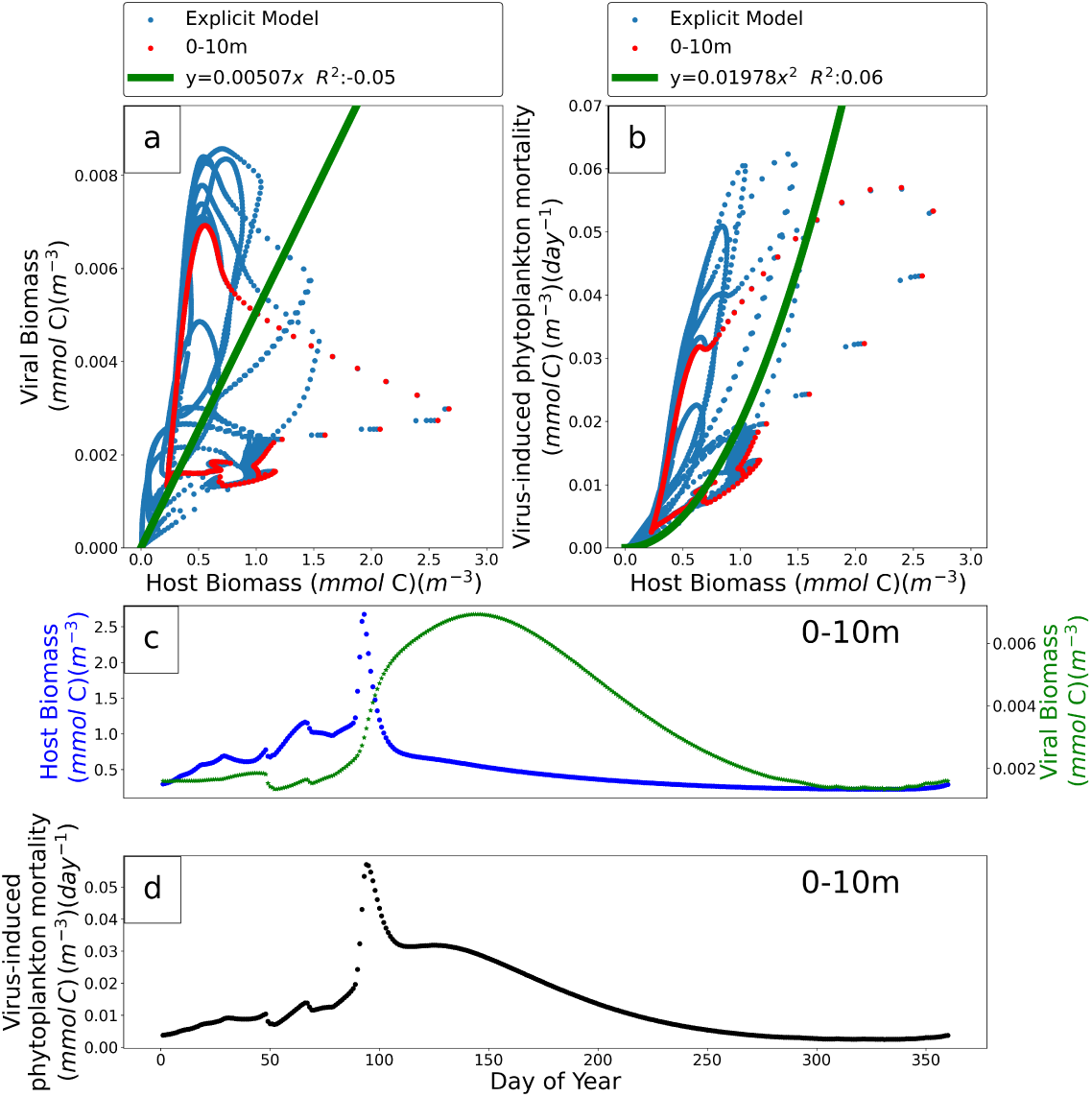
Comparison between implicit and explicit representations of viral-induced phytoplankton mortality in a one-dimensional configuration of the vDarwin-MITgcm system (see Methods and Frémont et al. (2026b) for details). Shown are the relationships between virus biomass density (a) and virus-induced mortality (b) with host biomass density. In both cases, the explicit model solutions are represented with blue dots, and the green lines represent the theoretical relationships that are required for the implicit assumption to be valid (see Figure 1b). The implicit model captures less than 10% of the variance of the explicit model relationship between biomass and virus-induced phytoplankton mortality. c) temporal dynamics of host biomass (blue) and viral biomass (green) for the top 0-10m. The same values are marked by the red dots in (a,b). d) temporal dynamics of virus-induced mortality in the top 0-10m. Model parameters are in Table 1.

Uncorrelated host and biomass densities are associated with temporal dynamics reminiscent of predator-prey cycles (Figure 2c), which can be dampened by a range of known ecological mechanisms (Steele & Henderson, 1992; Edwards, 2000; Omta et al., 2023). Arguably the simplest dampening processs is a density-dependent, quadratic loss on the virus which is included in the explicit model, and appears as ‘higher-order losses’ in Equation 5. Frémont et al. (2026a) showed that, when solved with static environmental forcing in a non-spatial environment, the Equations and parameters assumed in the explicit model lead the system to converge upon equilibrium states via damped oscillations. This cyclic behavior may be promoted within the 1-D vDarwin-MITgcm system as spatio-temporal variability arising from seasonality in light and nutrient availability, as well as advective and diffusive transport, may continually perturb the system and prevent it from reaching its equilibrium state. As a result of the dynamics and feedbacks inherent to the vDarwin-MITgcm system, temporal dynamics of virus-induced mortality are non-linear (Figure 2d), and cannot be captured implicitly.

### 3.2 Capturing ecosystem properties in a one-dimensional water column model

We next assumed the fitted value of the quadratic mortality coefficient in Figure 2 (*δ_vv_*=0.01978 m^3^ mmol*^−^*^1^ day*^−^*^1^) to evaluate the ability of the implicit model to capture the effects on ecosystem properties such as plankton biomass density and primary production that were evident within the explicit model. In Figure 3 we show depth-profiles of plankton biomass, virus-induced mortality, and primary production across two seasonal cycles for the explicit (left-column) and implicit (right-column) models. In both cases, environmental forcing and vertical transport are configured to match the Bermuda Atlantic Time-Series site (Wu et al., 2021), and the only difference is the representation of virus-induced mortality in Equation 2. In both the explicit and implicit representations, the phytoplankton biomass is low in January through to March, but begins to increase in April and onward before peaking at the onset of stratification in May. Virus-induced mortality and zooplankton biomass both increase in response to increases in rising phytoplankton biomass, but the increase is sustained until much later in the summer when virus-induced mortality is explicitly resolved. The implicit model shows a more abrupt seasonal decline in virus-induced mortality. Although zooplankton biomass and primary production follow qualitatively similar patterns in the explicit and implicit case, the zooplankton biomass density is slightly higher during the stratified summer months when the virus-induced mortality is implicit. This difference points to ecosystem wide implications of the representation of virus-induced mortality.

**Figure 3:**
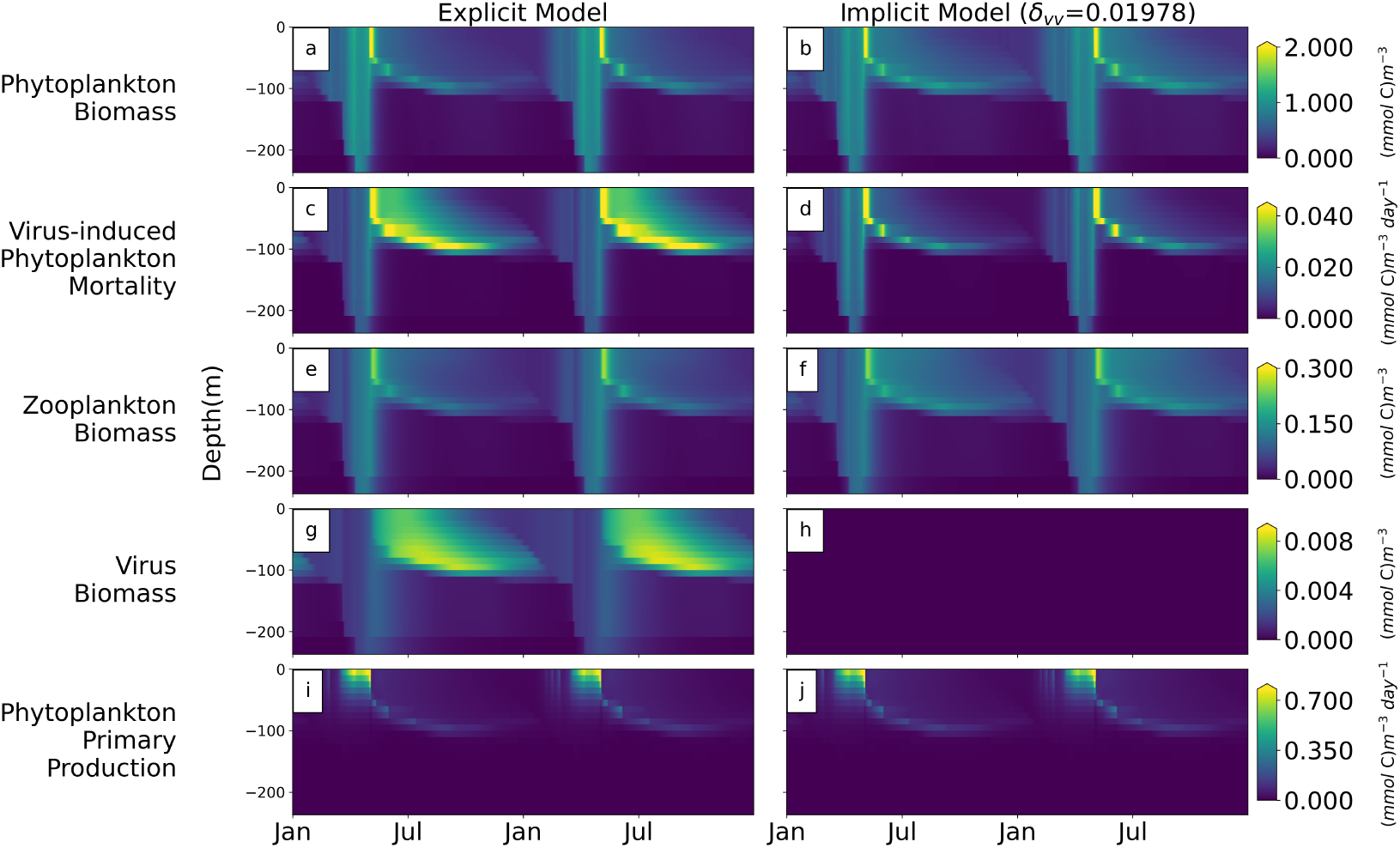
Comparison of ecosystem properties in the explicit (left-column) and implicit (right-column) versions of vDarwin within a one-dimensional water-column configuration of the MITgcm consistent with the Bermuda Atlantic Time-Series site (BATS) (Dutkiewicz et al., 2001). The implicit model assumes the quadratic mortality coefficient determined by fitting the implicit representation of virus-induced mortality to the explicit model (Figure 2b). Two modeled years are shown to demonstrate the repeating seasonal trend. Patterns in plankton biomass and primary production are qualitatively similar in the explicit and implicit model, but virus-induced mortality in the explicit model is sustained well into the summer months once the water column has become stratified.

The implicit model assumes any variation in virus-induced mortality occurs instantaneously in response to changes in host biomass density, whereas the explicit representation of viral dynamics allows a lag associated with the time taken for the viruses to replicate to sufficient numbers before they can meaningfully drive host biomass downward. These differences are evident in the second row of Figure 4, where red regions indicate the explicit model overpredicts the implicit model dynamics for phytoplankton biomass and virus-induced mortality, and blue regions indicate the opposite. The vertical red band spanning the top 200m in the left-column, middle row of Figure 4 highlights a period where phytoplankton biomass is beginning to increase at the onset of spring, where the explicit model predicts greater biomass than the implicit model. Here, viruses are yet to accumulate to significant numbers in the explicit model (Figure 3), and virus-induced mortality of phytoplankton is minimal. There is a delay in the rise in viral biomass in the explicit model, but once the viruses (and phytoplankton) reach sufficient numbers, the explicit model shows reduced phytoplankton biomass by comparison to the implicit model, marked by the shallow blue band that persists throughout the summer and into the autumn. Reduced phytoplankton biomass in the explicit model explains why the zooplankton have slightly lower biomass during stratification in the explicit case (compare Figure 3e,f), as their prey become less abundant. When viruses are modeled explicitly, the delay between infection and lysis allows the virus to diminish the biomass the phytoplankton accumulate as blooms progress, and these effects cascade through trophic levels, to the zooplankton.

**Figure 4:**
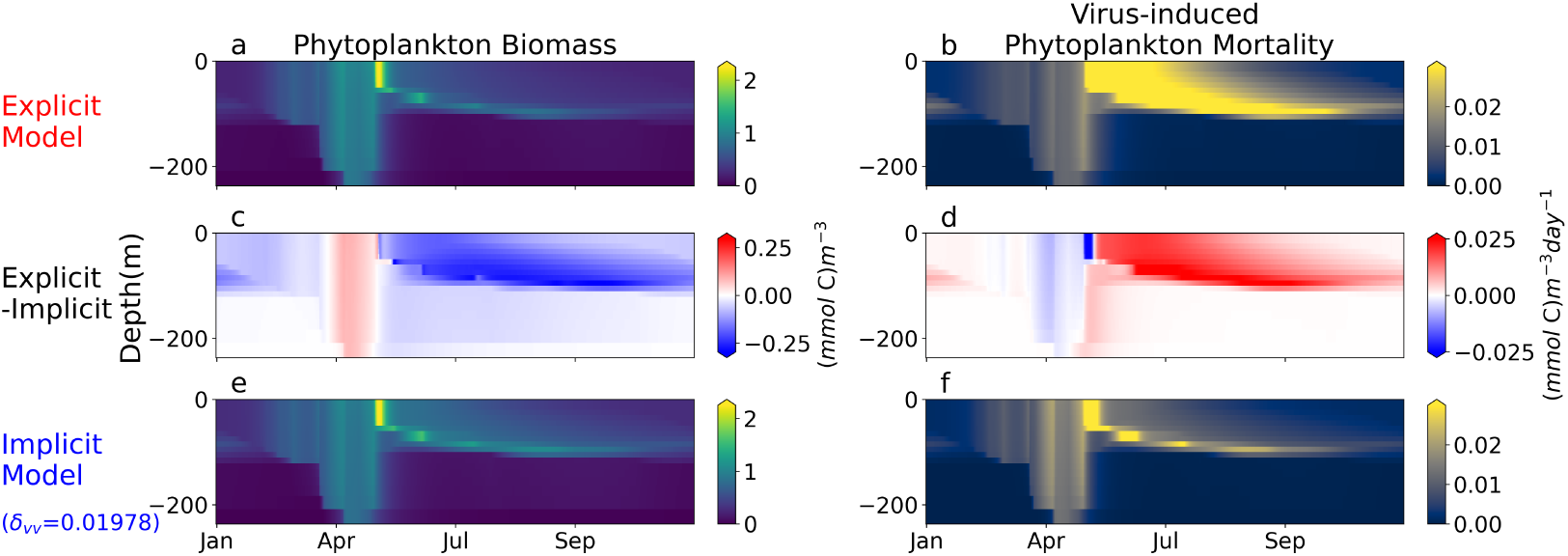
Comparison of spatio-temporal patterns of phytoplankton biomass and virus-induced mortality for explicit and implicit representations of virus-induced mortality. The middle row subtracts results of the implicit model (bottom-row) from the explicit model (top-row), with red showing over-prediction by the explicit model, and blue showing the opposite. The implicit model stymies phytoplankton biomass accumulation in the spring (vertical red band in April-May) whereas the explicit model predicts more severe viral impacts much later in the year once the viruses have had time to replicate.

The corresponding representation of virus-induced mortality in the right-hand column of Figure 4 shows the opposite pattern - a vertical blue band in the spring turns red in the summer, when viruses and phytoplankton have both accumulated to sufficient density that their contact rates sustain significant virus-induced mortality. The early control imposed on phytoplankton biomass by the implicit model stymies phytoplankton biomass accumulation by comparison to the explicit case. Interestingly, there is a brief period at the onset of stratification where the dominant representation becomes depth-dependent (Figure 4d where blue changes to red with depth). Here, the phytoplankton biomass is at its peak at the surface in both models (Figure 4a,e), which shades the phytoplankton residing deeper in the water-column. The temporal shift is the same at the surface and beneath 200m, but the reduced biomass at depth diminishes the amplitude of the differences (shown by fainter coloring at depth) and leads to truncated dynamics.

The comparisons in Figures 3,4 reveal considerable spatio-temporal variability in the dynamics of implicit and explicit representations of virus-induced mortality, and suggest that the implicit model is unable to adequately represent viral influences on marine ecosystem dynamics. We next asked whether values of the quadratic mortality coefficient other than the one derived by fitting the relationship between virus-induced mortality and host biomass in the explicit model (Figure 2) led to closer agreement between implicit and explicit representations of host biomass, virus-induced mortality, and primary productivity (Figure 5). Different values of *δ_vv_* must be assumed to minimize agreement between biomass, virus-induced mortality, and primary production, and none of these are consistent with the value of *δ_vv_* that best explains the relationship between virus-induced mortality and host biomass (Figure 2) (compare red and cyan lines in Figure 5). These differences reflect the non-linear structure of the vDarwin-MITgcm system, and highlight inherent trade-offs that arise when implicitly representing viral effects on marine ecosystems.

**Figure 5:**
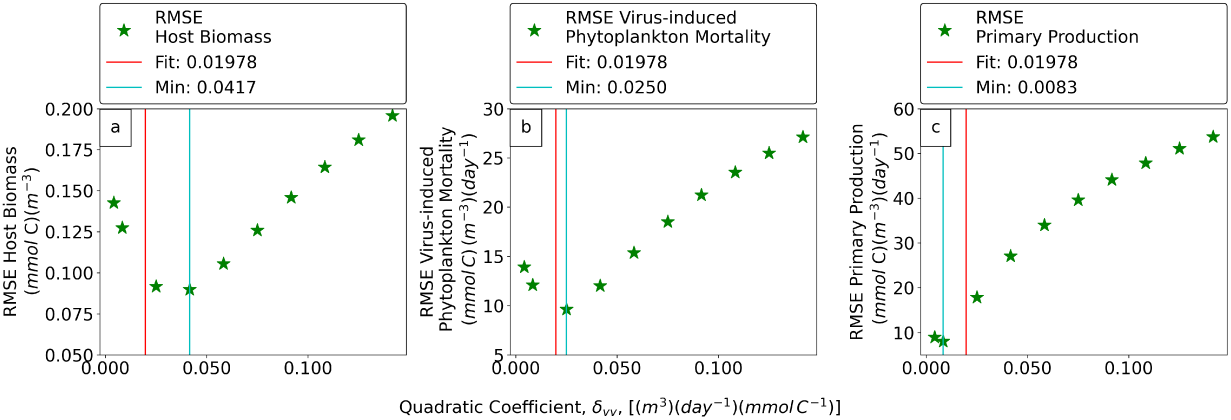
Sensitivity analysis examining the difference between explicit and implicit model predictions of simulated phytoplankton biomass (a), virus-induced mortality (b) and primary production (c) for many different values of the quadratic mortality coefficient *δ_vv_*. The vertical red line across panels mark the value of the coefficient assumed in Figures 2-4, and the cyan lines mark the value of the coefficient that minimized the root mean squared error (RMSE) between the implicit and explicit models when a range of different values for the coefficient (marked by green stars) were assumed in one-dimensional simulations of vDarwin-MITgcm. RMSE’s were calculated using 1-day averaged model output. Quadratic coefficients were evenly sampled within the x-axis range, with one additional simulation to verify that the RMSE for primary production reaches a minimum close to the origin.

### 3.3 Global ocean simulations

We next evaluated differences between explicit and implicit representations of viral infection in a three-dimensional global-ocean configuration of the vDarwin-MITgcm system. The left and right columns in Figure 6 show annually averaged, depth integrated simulations for the explicit and implicit case, respectively, and the middle column shows normalized differences between the two, where the differences were calculated prior to averaging. The value of the quadratic coefficient (*δ_vv_*=0.125) was determined via sensitivity analyses described below. The patterns in phytoplankton biomass and productivity in the left and right columns of Figure 6 are qualitatively consistent with expectations from ocean color (McClain, 2009) and primary production models (Carr et al., 2006), reflecting increases in nutrient availability in equatorial upwelling and coastal environments, and seasonally stratified regions at high latitude, relative to permanently stratified oligotrophic regions. The middle column shows significant spatial heterogeneity in the comparison between the implicit and explicit model, suggesting the implicit model does not recapitulate the influence of viral infection on marine ecosystems.

**Figure 6:**
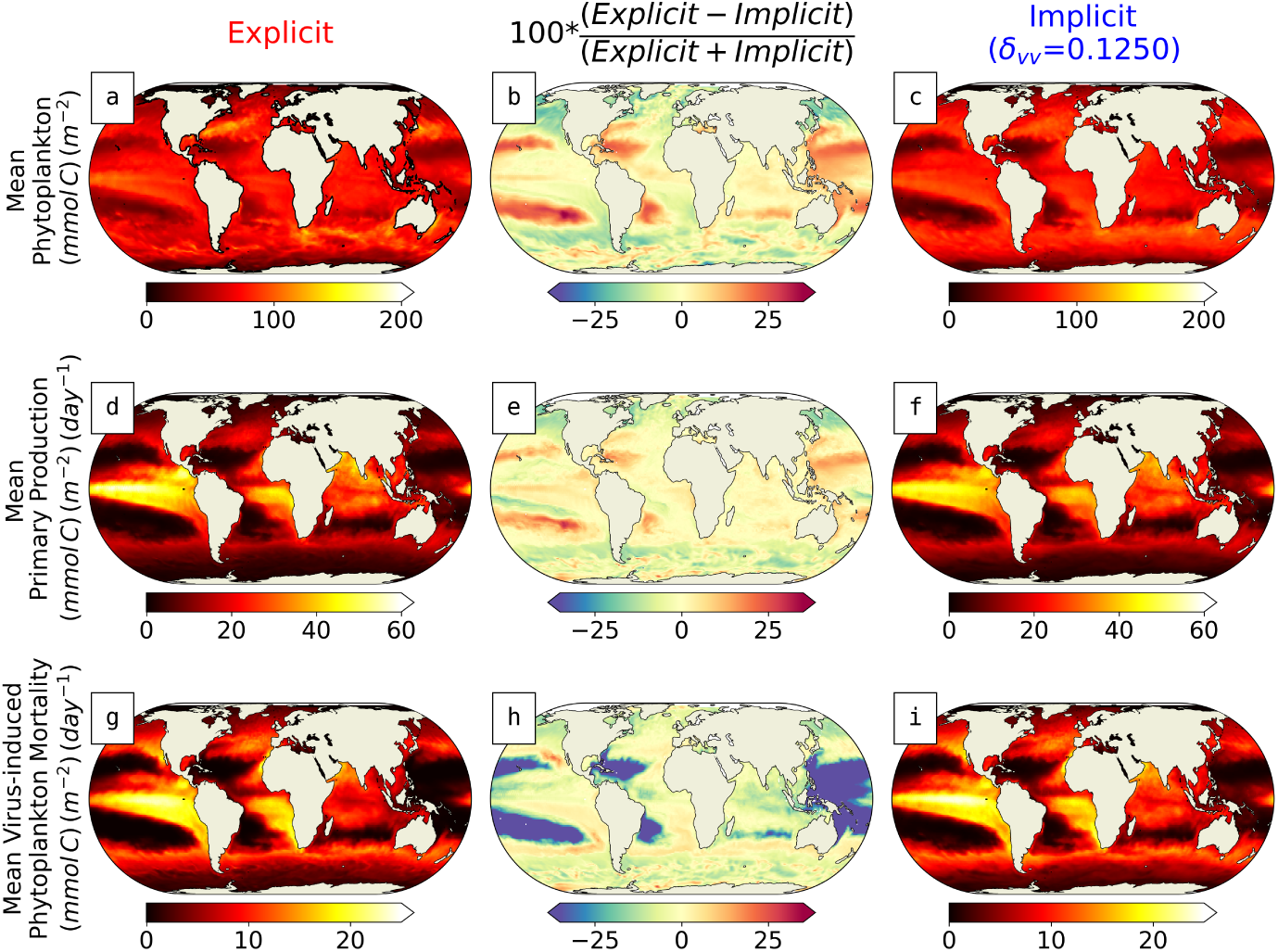
Global ocean simulations comparing explicit and implicit representations of viral infection within the vDarwin-MITgcm system. Shown are annually averaged, depth integrated simulations of phytoplankton biomass (top-row) primary production (middle-row) and virus-induced mortality (bottom-row). Simulations with explicit and implicit viral infection are shown in the left and right columns, respectively, and the middle column shows differences between the two, with red reflecting over-prediction by the explicit model, and blue the opposite. The spatial variation evident in the middle column suggests implicit representations of viral infection do not recapitulate viral influences on marine ecosystems.

In the least productive oligotrophic regions, the implicit model leads to lower predictions of phytoplankton biomass density and primary production by comparison to the explicit model. This distinction is marked by the blue regions at low latitude in Figure 6b,e and correspond to areas where the implicit model of viral infection predicts relatively high rates of virus-induced mortality. Here, when viruses are modeled explicitly, they must sustain sufficient replication to overcome losses associated with decay and particle attachment (Equation 5). Low host biomass in oligotrophic regions limits host-virus contact, preventing viruses from imposing significant mortality. The need for viruses to overcome their own losses is not resolved in the implicit model, where some degree of virus-induced mortality is assumed whenever host biomass is present.

As the system shifts toward higher productivity the pattern becomes more complicated, with a more general tendency of the implicit model to over-predict biomass, production, and virus-induced mortality. This pattern is reminiscent of the dynamics that emerge after bloom-formation in one-dimensional simulations (Figures 3,4), pointing to a failing of the implicit model to capture the ability of viruses to constrain phytoplankton blooms. This limitation is most pronounced at high latitudes, where phytoplankton biomass is far more dynamic seasonally. Here, the implicit model predicts large biomass inventories by comparison to the explicit model. These results suggest that when viruses are explicitly resolved, they have the propensity to act as strong ‘bloom terminators’ even when the nutrients are clearly available to support higher phytoplankton biomass density. These differences become even more pronounced when a lower value for the quadratic mortality coefficient is assumed (Figure A1), highlighting the sensitivity of the system to this parameter. The high latitudes where this pattern is most evident have relatively deep mixed layers and accumulate far higher biomass than the BATS site simulated in Figures 3 and 4, and resulting high rates of host-virus contact may drive rates of virus-induced mortality that the phytoplankton are unable to overcome. The analysis points to significant limitations in the ability of the implicit model to capture impacts of viruses on phytoplankton biomass dynamics and primary production across broad spatial scales.

The global ocean simulations reported in Figure 6 assumed a value of the quadratic mortality coefficient that was determined via sensitivity analyses following the approach taken in the 1D simulations. As with the one-dimensional simulations, we began by comparing the relationships between virus biomass, host biomass, and virus-induced phytoplankton mortality with the theoretical relationships that are required for the implicit, quadratic assumption to hold (Figure 7). Here, the relationship between virus and host biomass is highly dynamic (orange hexagons, Figure 7a) and a linear model (green line) explains ∼ 40% of the variance. Similar patterns emerge in the relationship between virus-induced mortality and host biomass (Figure 7b), and the quadratic model (green line) explains ∼34% of the variance.

**Figure 7:**
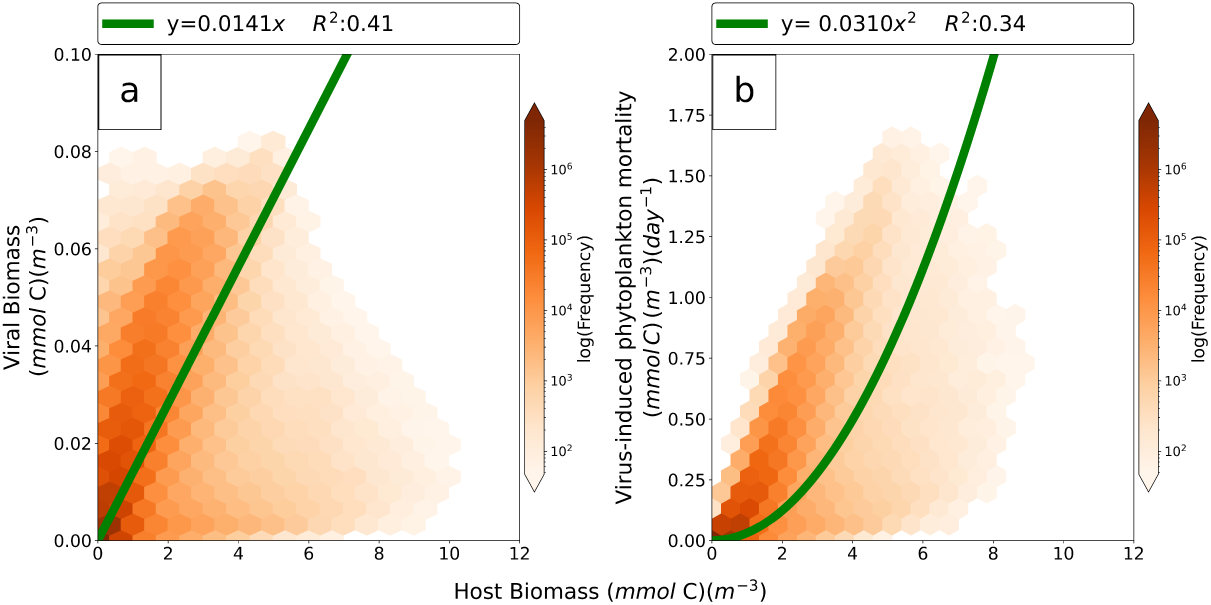
Comparison between implicit and explicit representations of virus-induced phytoplankton mortality in a three-dimensional configuration of the vDarwin-MITgcm system (see Methods and Frémont et al. (2026b) for details). Model parameters are in Table 1. Shown are the relationships between virus biomass density (a) and virus-induced mortality (b) with host biomass density. In both cases, the shade of orange represents the frequency of global model grid cells falling within the hexagons, and the green lines represent the theoretical relationships that are required for the implicit assumption to be valid (see Figure 1b). Model output was averaged over ten days prior to fitting. The implicit model captures less than 40% of the variance of the explicit model relationship between biomass and virus-induced phytoplankton mortality.

We next examined whether three-dimensional simulations warranted a different value of the quadratic mortality coefficient *δ_vv_* than the one we derived by fitting the quadratic function to the virus-induced mortality vs. biomass pattern in Figure 7b. In Figure 8 we show the root mean squared error (RMSE) between the explicit and implicit modeled representation of phytoplankton biomass (Figure 8a), virus-induced mortality (Figure 8b) and primary production (Figure 8c) for many values of the quadratic coefficient, *δ_vv_*. The value of *δ_vv_* that minimized the RMSE is marked by the vertical blue line in each panel, and is much higher than the one obtained in Figure 2b marked by the vertical red lines in each panel, as well as the values that were optimal for the one-dimensional simulations (Figures 2-5). These difference indicate that the implicit representation requires multiple, different values of the quadratic coefficient *δ_vv_* to capture the impact of virus-induced mortality on ecosystem properties. Note that in Figure 6, we assumed a value of *δ_vv_* that minimized the RMSE in Figure 8b (orange circle), as we anticipate this gives the implicit model the strongest possible representation of the influence of viruses on plankton biomass and primary production in this spatial setting. Equivalent simulations assuming the lower value optimal for curve fitting (marked by the red lines in Figure 8) are reported in Figure A1.

**Figure 8:**
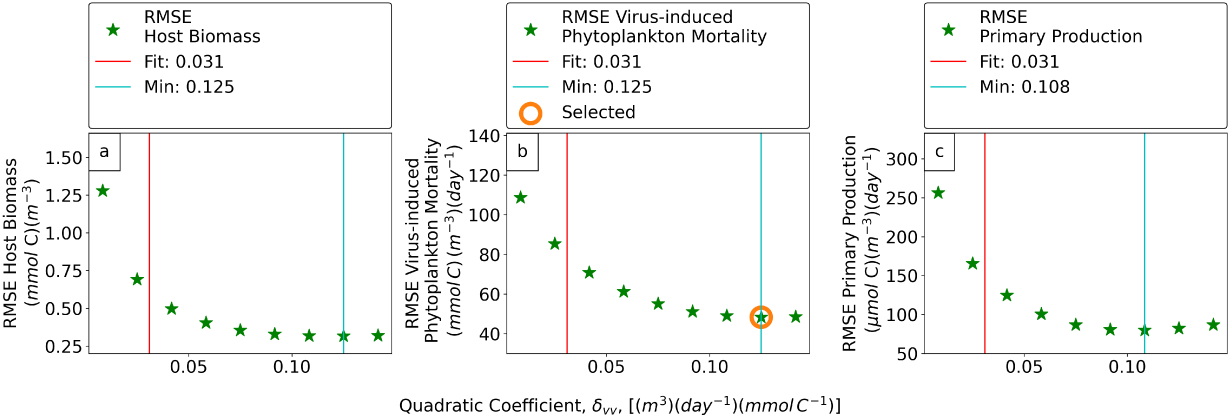
Sensitivity analysis examining the difference between explicit and implicit model predictions of simulated phytoplankton biomass (a), virus-induced mortality (b) and primary production (c) for many different values of the quadratic mortality coefficient *δ_vv_*. The red vertical line in each case is the quadratic coefficient that was fitted to the relationship between virus-induced mortality and biomass in Figure 7b, and the cyan lines mark the values that minimized the root mean squared error (RMSE) between the implicit and explicit model for each variable. RMSE’s were calculated using 10-day averaged model output and quadratic coefficients were evenly sampled within the x-axis range. The orange ‘selected’ circle marks the value of the quadratic coefficient assumed in Figure 6.

## 4 Discussion

Through cell lysis, viruses move organic carbon from hosts to detrital material (Suttle, 2007; Lara et al., 2017). Whether lysed material is recycled or retained within particulate and dissolved organic matter has implications for ocean biogeochemistry and ultimately the Earth system. Simulating the impacts of viral lysis in marine ecosystem models is a step toward assessing their effects on biogeochemical dynamics, and invites the question: Is it possible that viral effects can be captured implicitly, as others have assumed (Stock et al., 2014; Behrenfeld & Boss, 2014)? Here, we have addressed this question with a simple nutrient-phytoplankton-zooplankton-virus-detritus (NPZVD) model that lacks the diversity present in nature, but uses realistic life-history traits and simulates infection dynamics consistent with observations (Frémont et al., 2026b).

Our analyses reveal considerable spatio-temporal discrepancies between the different descriptions of viral infection. One-dimensional simulations indicate the implicit representation cannot capture the loss of phytoplankton biomass that occurs after stratification and viral biomass has accumulated during bloom formation. Three-dimensional simulations reveal broader spatial differences between the descriptions. The implicit representation contains no description of the losses viruses themselves must overcome - such as abiotic degradation and consumption by protists - which in our simulations with explicit viruses diminishes their ability to deplete phytoplankton biomass in low-resource environments, especially in the subtropical gyres. By contrast, the implicit model fails to capture the roles viruses play in phytoplankton bloom termination, especially at higher latitudes.

A key feature of viral infection is the time required for viruses to replicate to sufficient numbers to cause appreciable host mortality. This feature is challenging to capture without explicitly resolving the production and loss of viruses. Inclusion of these processes leads to nonlinear and uncorrelated host and virus biomass dynamics. This deviation from linearity is a direct violation of the assumption required for the quadratic assumption to hold (Figure 1b). We therefore conclude that the implicit representation of viral infection is unable to simulate the dynamics of virus-induced mortality that emerge when viruses are explicitly resolved. Furthermore, the implicit representation fails to capture the imprint of viral dynamics on key ecosystem features, such as primary production and phytoplankton biomass.

The NPZVD configuration of vDarwin assumed here was parameterized to simulate *Prochlorococcus* and its cyanophages. It is therefore most applicable to the oligotrophic regions where *Prochlorococcus* is numerically dominant (Flombaum et al., 2013). While caution should be applied interpreting results of the 3D model in more nutrient rich regions where eukaryotes tend to dominate (Ward et al., 2014), some lessons may nevertheless be gleaned. The representation of viral replication and extracellular loss assumed in the explicit model captures basic features common to all lytic viruses - their replication is contingent upon contact with their hosts, and their loss is due to particle attachment and decay. Extending vDarwin to capture the dynamics of viruses of eukaryotes (primarily pico-eukaryotes, coccolithophores, diatoms, and dinoflagellates) is a major endeavor, requiring development of appropriate molecular techniques to quantify rates of *in situ* infection along with robust quantification of life-history traits across diverse strains and ecotypes. Early efforts to characterize infection dynamics in these eukaryote-virus systems suggest the potential for the proportion of cells infected to be far higher than the ∼1% of hosts infected in the current version of vDarwin (Proctor & Fuhrman, 1990), which is an underestimate even for *Prochlorococcus* in regions beyond the North Pacific Subtropical Gyre (Carlson et al., 2022). It is therefore possible that future, more ecologically realistic vDarwin configurations will lead to greater divergence between explicit and implicit representations of virus-induced mortality.

The decision to incorporate explicit viral dynamics within models will depend upon the degree to which viral processes can reliably be constrained, and whether the uncertainty introduced by representation of viral infection is a reasonable payoff for the explanatory power it provides. The uncertainty associated, for example, with heterogeneity in life-history traits (Hinson et al., 2023; Edwards, 2018) will introduce uncertainty into models. Nevertheless, explicit representation of viruses within biogeochemical models is an imperative for Earth System, ecological, and biogeochemical modelers wishing to parameterize their effects. In our investigation, the value of the higher-order loss rate for viruses, *δ_vv_*, that minimized error between the implicit and explicit models could only be reliably estimated once both the implicit and explicit models were solved (Figure 8). The numerical value of *δ_vv_* that provides the optimal representation of viral dynamics is likely to change as models become more taxonomically representative. We therefore refrain from providing concrete recommendations for representation of viral dynamics within models until more ecologically realistic simulations have been developed, but advocate for expanded representation of viral dynamics within ecosystem and biogeochemical models to inform the representation of viral impacts in models more broadly.

The narrow taxonomic resolution of the NPZVD model used here is just one limitation preventing broader representation of viral dynamics in ecosystem models, and expansion to represent more groups will come with its own set of challenges. For example, infected cells were assumed to lyse their hosts at a constant rate. In reality, the timing of lysis in infected cells is subject to a range of environmental and molecular signals that are not well understood (Silveira et al., 2021). Furthermore, there is debate about the extent to which viruses shape community composition (Castledine & Buckling, 2024). The kill-the-winner hypothesis poses that viruses shape diversity by controlling otherwise dominant organisms (Thingstad, 2000; Winter et al., 2010). Ecoevolutionary dynamic models of viruses and their microbial hosts reveal mechanisms by which virus-induced mortality can drive the selection for expanded diversity, both of viruses and of hosts (Weitz et al., 2005; Beckett & Williams, 2013). An alternative view is that viruses play a more passive role, evolving the machinery to infect specific hosts whose prevalence and activity is shaped primarily by other environmental and ecological constraints (Castledine & Buckling, 2024). By including viruses in large-scale models we can begin to tease out the relevance of virus-induced effects beyond microbial diversity into the realm of biogeography, community structure, ecosystem function, biogeochemical cycles and more. We advocate for expansion of realism within models through continued quantification of molecular signatures of infection linked with life-history traits in ecologically representative model systems. There has been considerable progress quantifying molecular markers of infection in systems with well-constrained life-history parameters (Vardi et al., 2009; Vincent et al., 2021), with the potential to facilitate expanded representation and evaluation of the predictive value in incorporating viruses within ecosystem and biogeochemical models.

## Open Research Section

Code and data necessary to reproduce model figures are available on Zenodo at:

https://doi.org/10.5281/zenodo.21461475

vDarwin code modifications are available at:

https://github.com/werdna-spatial/ImpExpVirus.git

## Conflict of Interest disclosure

The authors declare there are no conflicts of interest for this manuscript.

## Acknowledgments

This work is supported by grants from the Simons Foundation: no. 721231, SFI-LS-PROJECT-00011159 and MPS-SIP-00930382 to JSW, no. 549931 and SFI-LS-Project-00011157 to SD, SFI-LS-Project-00010539 to DT, no. 721254 and SFI-LS-Project-00011154 to DL and SFI-LS-Project-01157186 and CBIOMES-00827829 to CF. PF is supported by a post-doctoral fellowship from the Simons Collaboration on Ocean Processes and Ecology (SCOPE), funded by the Simons Foundation through grants awarded to JSW. SJB and JSW are investigators at the University of Maryland-Institute for Health Computing, which is supported by funding from Montgomery County, Maryland and The University of Maryland Strategic Partnership: MPowering the State, a formal collaboration between the University of Maryland, College Park and the University of Maryland, Baltimore. DT acknowledges NSF OCE grants no 2445508 and no 2023680.

## Author Contributions

Conceptualization: DT; methodology: DT, EC, SD, JSW; model development: DT, EC, PF, SJB, DM, DD, DM, EC, OJ, CLF, DT, DL, JSW, SD; writing—review & editing: PF, SJB, DM, DD, CLF, DT, DL, JSW, SD; funding acquisition: CLF, DT, DL, JSW, SD.

## Appendix A Supplementary global sensitivity result

**Figure A1:**
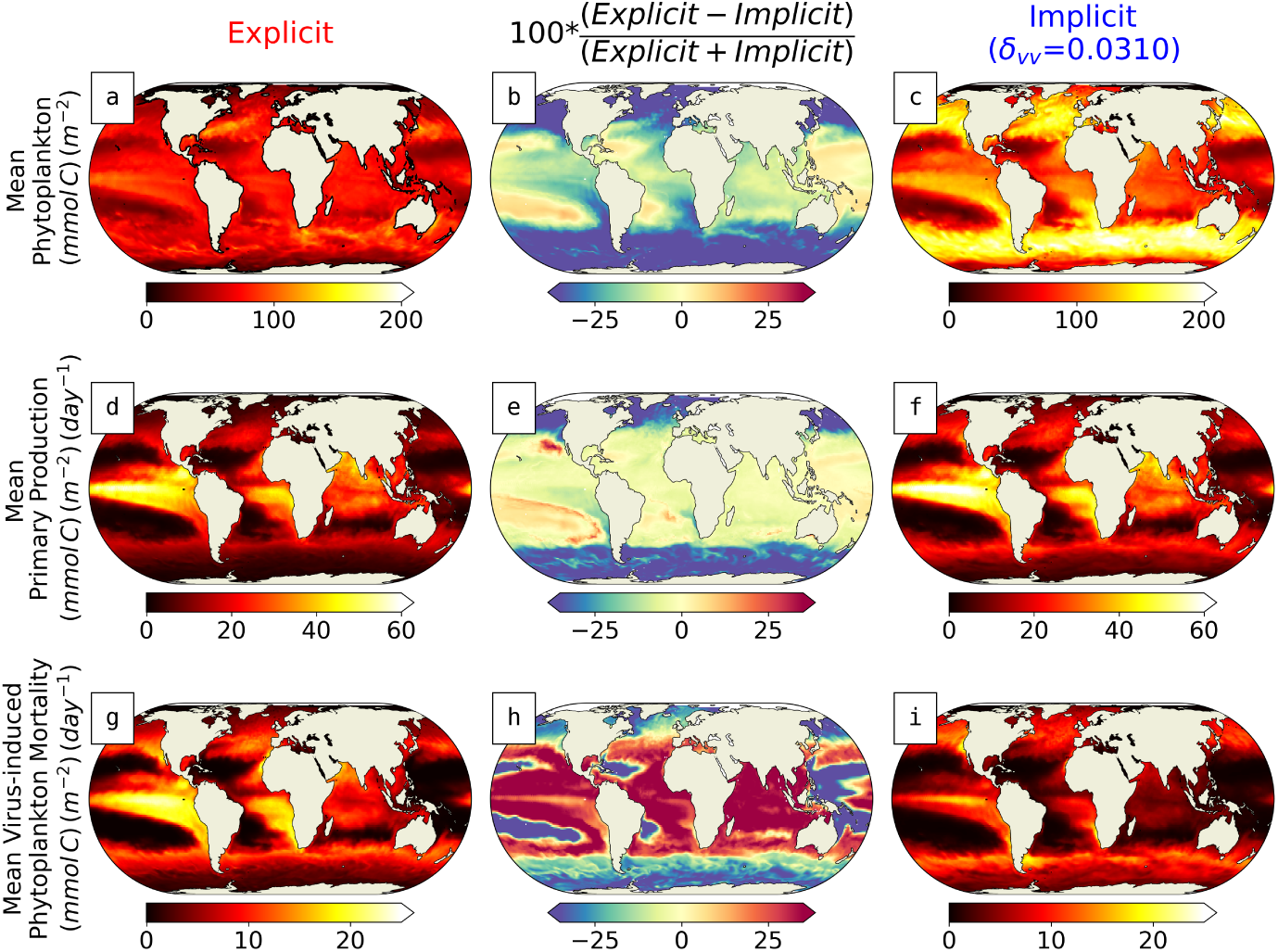
Global ocean simulations comparing explicit and implicit representations using the lower value of the quadratic coefficient (QC) obtained by fitting the quadratic function to relationship between virus-induced mortality and biomass in Figure 2. Shown are yearly averaged, depth integrated simulations of phytoplankton biomass (top-row) primary production (middle-row) and virus-induced mortality (bottom-row). Simulations with explicit and implicit viral infection are shown in the left and right columns, respectively, and the middle column shows differences between the two, with red reflecting over-prediction by the explicit model, and blue the opposite. The spatial variation evident in the middle column suggests implicit representations of viral infection do not recapitulate viral influences on marine ecosystems.

## Notes

### Competing Interest Statement

The authors have declared no competing interest.

